# Circulatory exosomes from COVID-19 patients trigger NLRP3 inflammasome in endothelial cells

**DOI:** 10.1101/2022.02.03.479081

**Authors:** Subhayan Sur, Robert Steele, T. Scott Isbell, Ranjit Ray, Ratna B. Ray

**Author notes:** **Address correspondence to** Ratna B. Ray, Department of Pathology, Saint Louis University, DRC, 207, 1100 South Grand Boulevard, St. Louis, MO 63104.

## Abstract

SARS-CoV-2 infection induces inflammatory response, cytokine storm, venous thromboembolism, coagulopathy, and multiple organ damage. Resting endothelial cells prevent coagulation, control blood flow and inhibit inflammation. However, it remains unknown how SARS-CoV-2 induces strong molecular signals in distant cells for immunopathogenesis. In this study, we examined the consequence of human endothelial cells (microvascular endothelial cells (HMEC-1) and liver endothelial cells (TMNK-1)) to exosomes from plasma of severe COVID-19 patients. We observed a significant induction of NLRP3, caspase-1 and IL-1β mRNA expression in the endothelial cells following exposure to exosomes from plasma of COVID-19 patients as compared to that of healthy donors. Activation of caspase-1 was noted in the endothelial cell culture medium following exposure to the COVID-19 exosomes. Further, COVID-19 exosomes significantly induced mature IL-1β secretion in the endothelial cell culture medium. Thus, our results demonstrated for the first time that exosomes from COVID-19 plasma trigger NLRP3 inflammasome in endothelial cells of distant organs.

## IMPORTANCE

SARS-CoV-2 infection is a global health problem. Although vaccine controls infection, understanding the molecular mechanism of pathogenesis will help in developing future therapies. Further, several investigators predicted the involvement of endothelial cell related inflammation in SARS-CoV-2 infection and using extracellular vesicles as a cargo to carry a drug or vaccine for combating against SARS-CoV-2 infection. However, the mechanism by which endothelial cells are inflamed remains unknown. Our present study highlights that exosomes from COVID-19 patients can enhance inflammasome activity in distant endothelial cells for augmentation of immunopathogenesis and opens an avenue in developing therapies.

SARS-CoV-2 infection, viral pathogenesis and resulting fatal multi-organ damage are global health concern. SARS-CoV-2 envelop spike protein interacts with angiotensin-converting enzyme 2, which is present on many cell surfaces as a receptor, but lung epithelial cells are probably the most susceptible cells for virus entry and replication causing human disease. Clinical observations indicate that severely ill COVID-19 patients experience chronic inflammation, cytokine storm, venous thromboembolic, coagulopathy and develop extra-pulmonary tissue/organ dysfunctions. The pathophysiology of these diverse manifestations is not clear but thought to occur in part to a dysregulated inflammatory response of immune cells and endothelial cells.

The innate immune system is the first line of defense system by which human body recognize and eliminate foreign pathogenic infection through involvement of highly conserved sensors, called pattern recognition receptors (PRRs). Inflammasomes, a high molecular weight cytoplasmic multiprotein complex of sensor protein and inflammatory caspase, play crucial role in sensing the external stimuli as well as in inducing cellular response against it. The NLR family pyrin domain containing 3 (NLRP3) is a type of PRR and most extensively studied inflammasome responsible for inflammation and antiviral responses (1). The NLRP3 inflammasome recruits pro-caspase-1 and activates caspase-1 by proteolytic cleavage resulting in caspase-1-dependent proteolytic maturation and secretion of interleukin (IL)-1β (2). A range of stimuli during pathogenic infections, tissue damage, or metabolic imbalances activate NF-kB which transactivates various effector genes including NLRP3, pro-IL-1β (1).

Aberrant activation of NLRP3 inflammasome or chronic inflammation triggered cellular damage resulting in severe pathological injury (1). Viral infection causes chronic inflammation and triggers NLRP3 inflammasome activation in immune cells. The hepatitis C virus including core protein, SARS-CoV viroporin, influenza virus M2, encephalomyocarditis virus viroporin 2B induces NLRP3 inflammasome activation (3–5). The non-immune cells such as epithelial cells, endothelial cells, and fibroblasts also contribute to innate immunity (6). Hyperinflammation, venous thromboembolic, dysregulated blood clotting and multiple organ damage in severely ill COVID-19 patients are suggested to dysfunction of endothelial cells (7). and need careful investigation.

Exosomes are extracellular vesicles (30–150 nm) and formed by the interior budding of endosomal membranes to form large multi-vesicular bodies. Exosomes play an important role in cellular communication and disease pathogenesis. Exosomes are also involved in viral spread, immune regulation and antiviral response during infection (8, 9). We have previously reported that exosomes released from hepatitis C virus (HCV) infected hepatocytes enhance fibrogenic markers in hepatic stellate cells (10). However, the functional role of exosomes from SARS-CoV-2 infected cells on macrophages and distant organs for pathogenic consequences remains unknown. Active NLRP3 inflammasome in PBMCs and tissues of postmortem patients upon autopsy was also reported (11), although supporting mechanism for this observation are not well defined. SARS-CoV-2 is not known to cause viremia, and clinical data indicate that virus infected individuals show other organ abnormalities during infection. SARS-CoV-2 directly or indirectly triggers inflammasomes, leading to the pleiotropic IL-1 family cytokines (IL-1β and IL-18) secretion (12), however, the molecular mechanisms for COVID-19 disease progression remain poorly understood.

We recently found that tenascin-C and fibrinogen-β are highly abundant in exosomes from COVID-19 patient plasma (13). Subsequently we showed that exosomes from COVID-19 patients trigger inflammatory signals to hepatocytes by inducing NF-kB through tenascin-C and fibrinogen-β. Thus, we hypothesized that exosomes from COVID-19 patients influence endothelial cell dysfunction and inflammatory response at distant organs. In this study, we found that exosomes from COVID-19 patients stimulates NLRP3 inflammasome formation and IL-1β production in endothelial cells.

## Results

### Exosomes from COVID-19 patients trigger inflammasome genes in endothelial cells

The COVID-19 patients were admitted to the ICU in our academic medical center, and the samples were collected on the date of arrival. The exosomes from plasma of healthy subjects (n=8) and COVID-19 patient (n=20) were isolated. The exosomes were characterized by expression of CD63 and TSG101 by western blot analysis, and further confirmed by transmission electron microscopy as described previously (13). To know the effect of COVID-19 exosomes on inflammasome formation in endothelial cells, human endothelial cells, HMEC-1 or TMNK1, were treated with normal or COVID-19 exosomes. We observed significant increase in mRNA expression of NLRP3, pro-caspase-1 (CASP1) and pro-IL-1β genes in the HMEC-1 cells exposed to exosomes from plasma of severe COVID-19 patients as compared that of health subjects (Figure 1A). Similar observation was noted with TMNK-1 cells following exposure to the COVID-19 exosomes (Figure 1B). This indicates that COVID-19 exosomes transcriptionally induce NLRP3 inflammasome components in endothelial cells of two different organs. We similarly incubated THP-1 cells (a human cell line derived from an acute monocytic leukemia patient, with or without PMA treatment) with exosomes from plasma of COVID-19 patients or that of healthy subjects, and a detectable change of NLRP3 inflammasome signaling was not observed.

**Figure 1:**
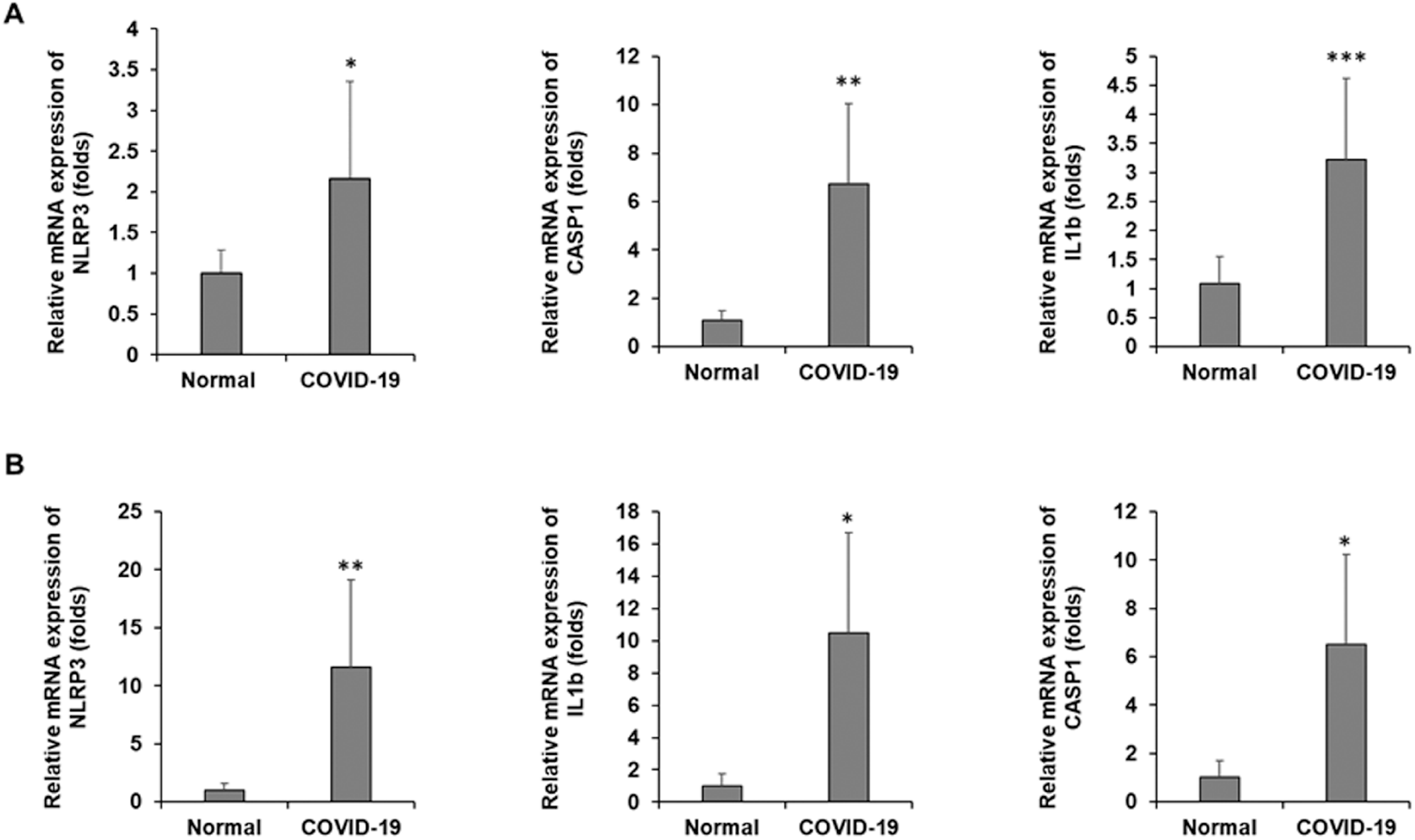
Exosomes from COVID-19 patients induce inflammasome genes. **A:** HMEC-1 and **B:** TMNK-1 cells were exposed to exosomes isolated from healthy subjects (n=8) and referred as normal or COVID-19 (n=20) patients for 48 h. Total RNA was isolated and relative mRNA expression of NLRP3, caspase-1 (CASP1) and 1L-1β were measured by qRT-PCR. 18s rRNA was used as an internal control. Small bar indicates standard error (* p<0.05; **, p<0.01; ***, p<0.001).

### Exosomes from COVID-19 patients activate caspase-1 in endothelial cells

The NLRP3 inflammasome is formed by oligomerization of NLRP3 proteins in cytoplasm (4). Functionally active inflammasome recruits and activates pro-caspase-1 by proteolytic cleavage. We examined caspase-1 expression in the endothelial cells following exposure to COVID-19 exosomes. Western blot analysis revealed a significant increase of cleaved caspase-1 in COVID-19 exosome exposed HMEC-1 cells as compared to normal (health subjects) exosomes exposed cells (Figure 2A). Caspase-1 is relatively stable and is released from inflammasome activated cells (14). We measured active caspase-1 in the culture media of exosomes exposed cells and a significantly high caspase-1 activity was detected (Figure 2B).

**Figure 2:**
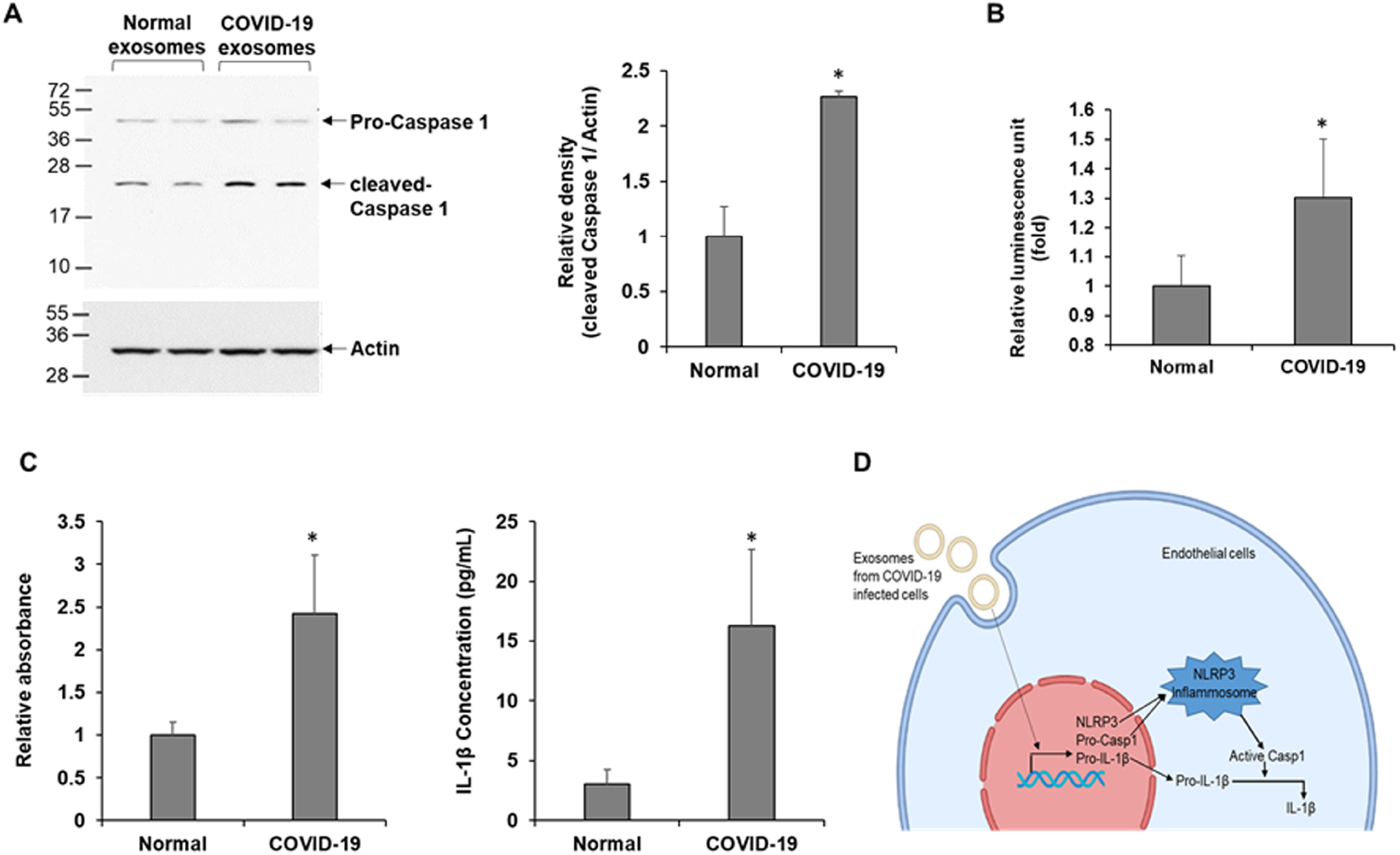
Exosomes isolated from COVID-19 patients activate caspase-1 and induce 1L-1β secretion. **A:** HMEC-1 cells were exposed to exosomes normal and COVID-19 patients for 48 h, and cell lysates were subjected to western blot analysis for caspase-1 using specific antibody. The membrane was reprobed for actin as an internal control. The quantitative presentation of band intensities using Image J software is shown. **B:** Caspase-1 activity was measured in exosomes exposed to HMEC-1 culture medium using Caspase-Glo 1 Inflammasome Assay reagent. Luminescence was read after 3 h incubation with the reagent and data presented as relative luminescence unit. **C:** HMEC-1 cells were exposed to exosomes from normal and COVID-19 patients for 48 h and IL-1β from culture medium was assayed using ELISA MAX Deluxe Set Human IL-1β kit. Relative absorbance was measured at 450 nm. Concentration of IL-1β in the media was calculated from the IL-1β standard curve. Small bar indicates standard error (*p<0.05). **D:** Schematic presentation shows exosomes secreted from SARS-CoV-2 infected cells trigger NLRP3, pro-caspase-1 (Casp1) and pro-IL-1β transcription resulting in activation of Casp1 followed by IL-1β via NLRP3 inflammasome in endothelial cells.

### Exosomes from COVID-19 patients induce maturation and secretion of IL-1β in endothelial cells

Active caspase-1 cleaves pro-IL-1β into its mature and functionally active cytokine IL-1 β. We examined mature IL-1β level in the culture media by ELISA. Exposure of COVID-19 exosomes significantly increased IL-1β level in culture media of HMEC-1 as compared to the normal exosome treated HMEC1 culture medium (Figure 2C). Similar increased IL-1β production was seen in the culture media of TMNK1 cells when exposed to COVID-19 exosomes (data not shown). Thus, COVID-19 exosomes appeared to trigger IL-1β production through activation of NLRP3 inflammasome in endothelial cells (Figure 2D).

## Discussion

Here, we investigated the effect of exosomes isolated from COVID-19 patients on endothelial cells of two distinct organs. Our study revealed that exosomes from COVID-19 patients trigger (i) NLRP3, caspase-1 and IL-1β transcription, (ii) NLRP3 inflammasome activation and (iii) maturation and secretion of IL-1β from endothelial cells. To our knowledge, this is the first demonstration of a mechanism for endothelial cell dysfunction by exosomes during COVID-19 disease.

Endothelial cells provide selectively permeable barrier to blood, regulate inflammation, platelet aggregation, thrombosis, and vascular smooth muscle proliferation (15). Resting endothelial cells prevent coagulation, control blood flow and passage of proteins from blood into tissues and inhibit inflammation. Endothelial cell dysfunction resulting in multi-organ failure is common feature of many viral infections including Influenza-A H1N1, SARS-CoV, MERS-CoV, Dengue virus (16, 17). Dengue virus NS1, NS2A and NS2B proteins also induce NLRP3 inflammasome, and IL-1β release in endothelial cells results in endothelial cell inflammation and dysfunction (17). Association of COVID-19 disease severity with pulmonary endothelial cell dysfunction, impaired microcirculatory function including venous thromboembolic disease and multiple organ involvement are reported (7). Deregulated host inflammatory response and cytokine storm are suggested to be driver of COVID-19 severity (18). Elevated level of IL-1β and its association with viral load and severity are also reported from COVID-19 patient blood (18, 19). However, effect of endothelial cell dysfunction in SARS-CoV-2 infection is not clearly known. A recent study demonstrated that SARS-CoV-2 N protein activates NLRP3 inflammasome and induces IL-1β and IL-6 production in epithelial cells and monocytes (20). However, we did not detect the SARS-CoV-2 N gene in COVID-19 patient exosomes. In proteomic analysis by mass-spectrometry, we did not detect NLRP3, caspase-1, IL-1β or NF-kB proteins in the COVID-19 patient exosomes (13). Although the mechanism of NLRP3 inflammasome activation is not known in endothelial cells, studies indicated that NF-kB, which is activated upon a range of stimuli during viral infection, transcribes effector genes NLRP3 and pro-IL-1β (1). We showed previously that tenascin-C and fibrinogen-β are highly abundant in exosomes from COVID-19 patients which activate NF-kB in hepatocytes (13), which may play a role in this process. We also observed SARS-CoV-2 spike protein, specially the S1 region, is present in the exosomes from plasma of patients. Although unmodified viral spike protein was widely used as a vaccine, diverse function of this protein was reported (21, 22). We therefore expressed SARS-CoV-2 spike protein in different cell lines and isolated the exosomes from culture media to incubate endothelial or THP1 cells. Interestingly we did not observe inflammasome formation in endothelial or THP1 cells (data not shown).

The mechanism of deregulation in multiple organs including, but not limited to, cardiac, neurologic, hemostatic, kidney, and liver during SARS-CoV-2 infection is not entirely clear. Exosomes play important role in cell-to-cell communication and viral pathogenesis. Many RNA viruses utilize the exosomal communication for viral pathogenesis (10, 23). For example, exosomes from hepatitis C virus infected hepatocytes carry materials for activation of hepatic stellate cells and induce fibrosis (10). Exosome mediated regulation of endothelial cells were reported in pregnant women (24). Exosomes from activated monocytes activates human brain microvascular endothelial cells to stimulate cytokines IL1β and IL6 through induction of NF-kB (25). Thus, exosomes may serve as an important mediator of endothelial cell dysfunction and inflammation in various organs during SARS-CoV-2 pathogenesis. In summary, our results suggested that COVID-19 plasma exosomes exposure induces NLRP3 inflammasome in endothelial cells of distant organs which may be one of the mechanisms of endothelial cell dysfunction and inflammation during severe COVID-19 disease.

The materials and Methods were provided as supplementary document.

## Acknowledgements

TMNK1 cells were kindly provided by A. Soto-Gutierrez, University of Pittsburg, Pittsburg, PA. This work was supported by the Research Institute of Saint Louis University (T.S.I. and R.R.) and Pathology Department Seed Grant to R.B.R.

## Authors’ contribution

RBR, TSI and RR conceived the research idea, SS, RS, RBR performed the experiments, SS, RR and RBR wrote the manuscript, and all authors edited and approved the final version of the manuscript.

